# Conserved and divergent aspects of Robo receptor signaling and regulation between *Drosophila* Robo1 and *C. elegans* SAX-3

**DOI:** 10.1101/2020.10.12.336297

**Authors:** Trent Daiber, Christine J. VanderZwan-Butler, Greg J. Bashaw, Timothy A. Evans

**Affiliations:** Department of Biological Sciences, University of Arkansas, Fayetteville, AR 72701; Department of Neuroscience, Perelman School of Medicine, University of Pennsylvania, Philadelphia, PA 19104

## Abstract

The evolutionarily conserved Roundabout (Robo) family of axon guidance receptors control midline crossing of axons in response to the midline repellant ligand Slit in bilaterian animals including insects, nematodes, and vertebrates. Despite this strong evolutionary conservation, it is unclear whether the signaling mechanism(s) downstream of Robo receptors are similarly conserved. To directly compare midline repulsive signaling in Robo family members from different species, here we use a transgenic approach to express the Robo family receptor SAX-3 from the nematode *Caenorhabditis elegans* in neurons of the fruit fly, *Drosophila melanogaster*. We examine SAX-3’s ability to repel *Drosophila* axons from the Slit-expressing midline in gain of function assays, and test SAX-3’s ability to substitute for *Drosophila* Robo1 during fly embryonic development in genetic rescue experiments. We show that *C. elegans* SAX-3 is properly translated and localized to neuronal axons when expressed in the *Drosophila* embryonic CNS, and that SAX-3 can signal midline repulsion in *Drosophila* embryonic neurons, although not as efficiently as *Drosophila* Robo1. Using a series of Robo1/SAX-3 chimeras, we show that the SAX-3 cytoplasmic domain can signal midline repulsion to the same extent as Robo1 when combined with the Robo1 ectodomain. We show that SAX-3 is not subject to endosomal sorting by the negative regulator Commissureless (Comm) in *Drosophila* neurons *in vivo*, and that peri-membrane and ectodomain sequences are both required for Comm sorting of *Drosophila* Robo1.

## Background

Members of the Roundabout (Robo) family of axon guidance receptors were identified in genetic screens in *Drosophila* and *C. elegans* by virtue of their mutant phenotypes, wherein subsets of axons exhibit guidance errors in homozygous mutant animals (Seeger *et al*. 1993; Zallen *et al*. 1999). The family is named after the *Drosophila roundabout (robo)* gene (later re-named *robo1)* and reflects that fact that axons in the embryonic CNS of *Drosophila robo1* mutants cross and re-cross the midline, forming aberrant circular pathways that resemble traffic roundabouts (Seeger *et al*. 1993). A screen for mutants exhibiting <u>s</u>ensory <u>a</u>xon (*sax*) defects in *C. elegans* identified mutations in a homologous gene, *sax-3/robo*, which also regulates midline crossing of axons in addition to its role in sensory axon guidance (Zallen *et al*. 1998; 1999). While *sax-3* is the only *robo* family gene in *C. elegans*, two additional *robo* genes were later identified in *Drosophila: robo2* and *robo3* (Rajagopalan *et al*. 2000a; Simpson *et al*. 2000b; Rajagopalan *et al*. 2000b; Simpson *et al*. 2000a). The three Robo receptors in *Drosophila* have distinct and in some cases overlapping axon guidance activities, but Robo1 is the main receptor for canonical Slit-dependent midline repulsion (Rajagopalan *et al*. 2000a; Simpson *et al*. 2000b).

Analyses of the Robo1 and SAX-3 protein sequences showed that these genes encode transmembrane receptor proteins with an evolutionarily conserved ectodomain structure including five immunoglobulin-like (Ig) domains and three fibronectin type III (Fn) repeats (Kidd *et al*. 1998a; Zallen *et al*. 1998). While there is little sequence similarity in the receptors’ cytoplasmic domains, three short conserved cytoplasmic (CC1, CC2, CC3) motifs were identified by virtue of their evolutionary conservation among Robo family members in flies, worms, and mammals (Kidd *et al*. 1998a). Later studies identified a fourth CC motif (named CC0) that was present in fly and human Robo proteins (Bashaw *et al*. 2000), but it was not clear whether CC0 was also present in the cytoplasmic domain of *C. elegans* SAX-3 (Simpson *et al*. 2000b; Dickson and Gilestro 2006).

Upon Slit binding, Robo receptors activate a cytoplasmic signaling pathway that induces collapse of the local actin cytoskeleton, resulting in growth cone repulsion. In *Drosophila*, midline repulsive signaling by Robo1 involves recruitment of downstream effectors that interact with the CC2 and CC3 motifs (Bashaw *et al*. 2000; Fan *et al*. 2003; Lundström *et al*. 2004; Hu *et al*. 2005; Yang and Bashaw 2006), as well as receptor proteolysis by the ADAM-10 metalloprotease Kuzbanian (Kuz) and clathrin-dependent endocytosis of proteolytically processed Robo1 (Coleman *et al*. 2010; Chance and Bashaw 2015). Phosphorylation of conserved tyrosine residues by the Abl tyrosine kinase (particularly within the CC1 motif) is important for negative regulation of Robo1-dependent signaling (Bashaw *et al*. 2000). *Drosophila* Robo1 is also subject to negative regulation by the endosomal sorting receptor Commissureless (Comm) (Tear *et al*. 1996; Kidd *et al*. 1998b; Keleman *et al*. 2002; 2005) and a second Robo family member, Robo2, which interacts with Robo1 in trans to inhibit premature Slit response in pre-crossing commissural axons (Simpson *et al*. 2000b; Spitzweck *et al*. 2010; Evans *et al*. 2015). As Comm and Robo2 appear to be conserved only within a subset of insect species, it is not yet clear how well conserved the signaling and regulatory mechanisms of *Drosophila* Robo1 are, although the mammalian Nedd4-family interacting proteins Ndfip1 and Ndfip2 can act analogously to *Drosophila* Comm to regulate surface levels of Robo1 on precrossing commissural axons in the spinal cord (Gorla *et al*. 2019) and the divergent Robo3/Rig-1 receptor in mammals appears to be able to antagonize Slit-Robo repulsion via a mechanism that is distinct from *Drosophila* Robo2 (Sabatier *et al*. 2004; Jaworski *et al*. 2010; Zelina *et al*. 2014). The mammalian PRRG4 protein can also regulate subcellular distribution of mammalian Robo1, similar to Comm’s effect on *Drosophila* Robo1, though whether PRRG4 influences midline crossing in mammalian neurons is not yet known (Justice *et al*. 2017).

Trans-species ligand-receptor binding experiments suggested that the mechanism of Slit-Robo interaction is highly conserved, as insect Slit can bind to mammalian Robos and vice-versa (Brose *et al*. 1999). Subsequent co-crystallization and biophysical studies of Slit-Robo interaction in insect and vertebrate Robos confirmed that the Slit-Robo interface between the Slit D2 domain and the Robo Ig1 domain were highly similar across species (Morlot *et al*. 2007; Fukuhara *et al*. 2008). Gain of function and genetic rescue experiments in *Drosophila* showed that Robo receptors from the flour beetle *Tribolium castaneum* can activate midline repulsion in fly neurons, even in the absence of *Drosophila robo1* (Evans and Bashaw 2012), while expression of human Robo1 (hRobo1) instead interfered with midline repulsion and produced dominant negative-like phenotypes in the fly embryonic CNS, suggesting that it is unable to signal repulsion in fly neurons (Justice *et al*. 2017).

Here, we examine the ability of SAX-3, the single *C. elegans* Robo ortholog, to signal midline repulsion in *Drosophila* embryonic neurons using gain of function and genetic rescue assays. We show that *C. elegans* SAX-3 can repel *Drosophila* axons from the Slit-expressing embryonic midline when misexpressed broadly or in specific subsets of commissural neurons, and can partially rescue ectopic midline crossing defects in *robo1* mutant embryos when expressed in *robo1’s* normal expression pattern. Using a panel of Robo1/SAX-3 chimeric receptors, we show that the cytoplasmic domain of SAX-3 can signal midline repulsion as effectively as the cytodomain of Robo1 when paired with the Robo1 ectodomain, indicating that the SAX-3 cytodomain can interact with and activate downstream repulsive signaling components in *Drosophila* neurons. We show that SAX-3 is not sensitive to endosomal sorting by the negative regulator Comm, which depends on sequences within the peri-membrane and ectodomain regions of Robo1, but SAX-3 is sensitive to sorting-independent Comm inhibition.

Together, our results provide evidence that repulsive signaling mechanisms of Robo family receptors are conserved outside of insects, and provide further insight into the structural basis for negative regulation of midline repulsive signaling by *Drosophila* Robo1.

## Materials and Methods

### Molecular Biology

#### pUAST cloning

The *sax-3* coding sequence was amplified via PCR from a *sax-3* cDNA template and cloned as an XbaI-KpnI fragment into a pUAST vector (p10UASTattB) including 10xUAS and an attB site for phiC31-directed site-specific integration. Robo1 and SAX-3 p10UASTattB constructs include identical heterologous 5’ UTR and signal sequences (derived from the Drosophila *wingless* gene) and an N-terminal 3xHA tag.

#### robo1 rescue construct cloning

Construction of the *robo1* genomic rescue construct was described previously (Brown *et al*. 2015). Full-length *robo1* and *sax-3* coding sequences were cloned as *BglII* fragments into the *BamHI*-digested backbone. Receptor proteins produced from this construct include the endogenous Robo1 signal peptide, and the 4xHA tag is inserted directly upstream of the Ig1 domain.

#### Robo1/SAX-3 chimeric receptors

*robo1* and *sax-3* receptor fragments were amplified separately via PCR, then assembled and cloned into the *robo1* rescue construct backbone using Gibson Assembly (New England Biolabs E2611). All coding regions were completely sequenced to ensure no other mutations were introduced. Robo1/SAX-3 variants include the following amino acid residues after the N-terminal 4xHA tag, relative to Genbank reference sequences AAF46887.1 (Robo1) and AAC38848.1 (SAX-3): robo1E-sax3PC (Robo1 Q52-Y890/SAX-3 N847-T1273); robo1EP-sax3C (Robo1 Q52-W973/SAX-3 N930-T1273); robo1EC-sax3P (Robo1 Q52-Y890/SAX-3 N847-Q929/Robo1 I974-T1395); sax3E-robo1PC (Robo1 Q52-S55/SAX-3 P31-M846/Robo1 H891-T1395); sax3EP-robo1C (Robo1 Q52-S55/SAX-3 P31-Q929/Robo1 I974-T1395); sax3EC-robo1P (Robo1 Q52-S55/SAX-3 P31-M846/Robo1 H891-W973/SAX-3 N930-T1273).

### Genetics

The following *Drosophila* mutant alleles were used: *robo1^1^* (also known as *robo^GA285^*) (Kidd *et al*. 1998a), *eg^Mz360^* (eg-GAL4) (Dittrich *et al*. 1997), comm^*E39*^ (Georgiou and Tear 2002). The following *Drosophila* transgenes were used: *P{GAL4-elav.L}3 (elavGAL4), P{10UAS-HARobo1}86Fb (UAS-Robo1), P{10UAS-HASAX3}86Fb (UAS-SAX3), P{UAS-CommHA}, P{robo1::HArobo1}* (Brown *et al*. 2015), *P{robo1:HAsax3}, P{robo1::HArobo1E-sax3PC}, P{robo1::HArobo1EP-sax3C}, P{robo1::HArobo1EC-sax3P}, P{robo1::HAsax3E-robo1PC}, P{robo1::HAsax3EP-robo1C}, P{robo1::HAsax3EC-robo1P}*. Transgenic flies were generated by BestGene Inc (Chino Hills, CA) using ΦC31-directed site-specific integration into attP landing sites at cytological position 86FB (for *UAS-SAX3*) or 28E7 (for *robo1* genomic rescue constructs). *robo1* rescue transgenes were introduced onto a *robo1^1^* chromosome via meiotic recombination, and the presence of the *robo1^1^* mutation was confirmed in all recombinant lines by DNA sequencing. All crosses were carried out at 25°C.

### Immunofluorescence and imaging

*Drosophila* embryo collection, fixation and antibody staining were carried out as previously described (Patel 1994). The following antibodies were used: FITC-conjugated goat anti-HRP (Jackson Immunoresearch #123-095-021, 1:100); mouse anti-Fasciclin II (Developmental Studies Hybridoma Bank [DSHB] #1D4, 1:100); mouse anti-βgal (DSHB #40-1a, 1:150); rabbit anti-GFP (Invitrogen #A11122, 1:1000); mouse anti-HA (Covance #MMS-101P-500, 1:1000); Cy3-conjugated goat anti-mouse (Jackson #115-165-003, 1:1000); Alexa 488–conjugated goat anti-rabbit (Jackson #111-545-003, 1:500). Embryos were genotyped using balancer chromosomes carrying *lacZ* markers, or by the presence of epitope-tagged transgenes. Ventral nerve cords from embryos of the desired genotype and developmental stage were dissected and mounted in 70% glycerol/PBS. Fluorescent confocal stacks were collected using a Leica SP5 confocal microscope and processed by Fiji/ImageJ (Schindelin *et al*. 2012) and Adobe Photoshop software.

## Results

### Sequence comparison of *C. elegans* SAX-3 and *Drosophila* Robo1

We compared the full-length protein sequences of *Drosophila* Robo1 and *C. elegans* SAX-3 to measure the degree of sequence similarity among the eight conserved ectodomain structural elements (5 Ig + 3 Fn) as well as the receptors’ cytoplasmic domains (Fig 1) (Kidd *et al*. 1998a; Zallen *et al*. 1998). The amino acid sequences of the eight ectodomain elements display varying degrees of evolutionary conservation, with 29-47% sequence identity between individual domains and the highest degree of sequence identity within Ig1 (46%), Ig3 (45%), and Ig5 (47%) (Fig 1A). Notably, the degree of sequence conservation within the Slit-binding Ig1 domain appears to be lower between *Drosophila* Robo1 and *C. elegans* SAX-3 (46% identical) than between *Drosophila* Robo1 and human Robo1 (58% identical) (Kidd *et al*. 1998a).

**Figure 1.**
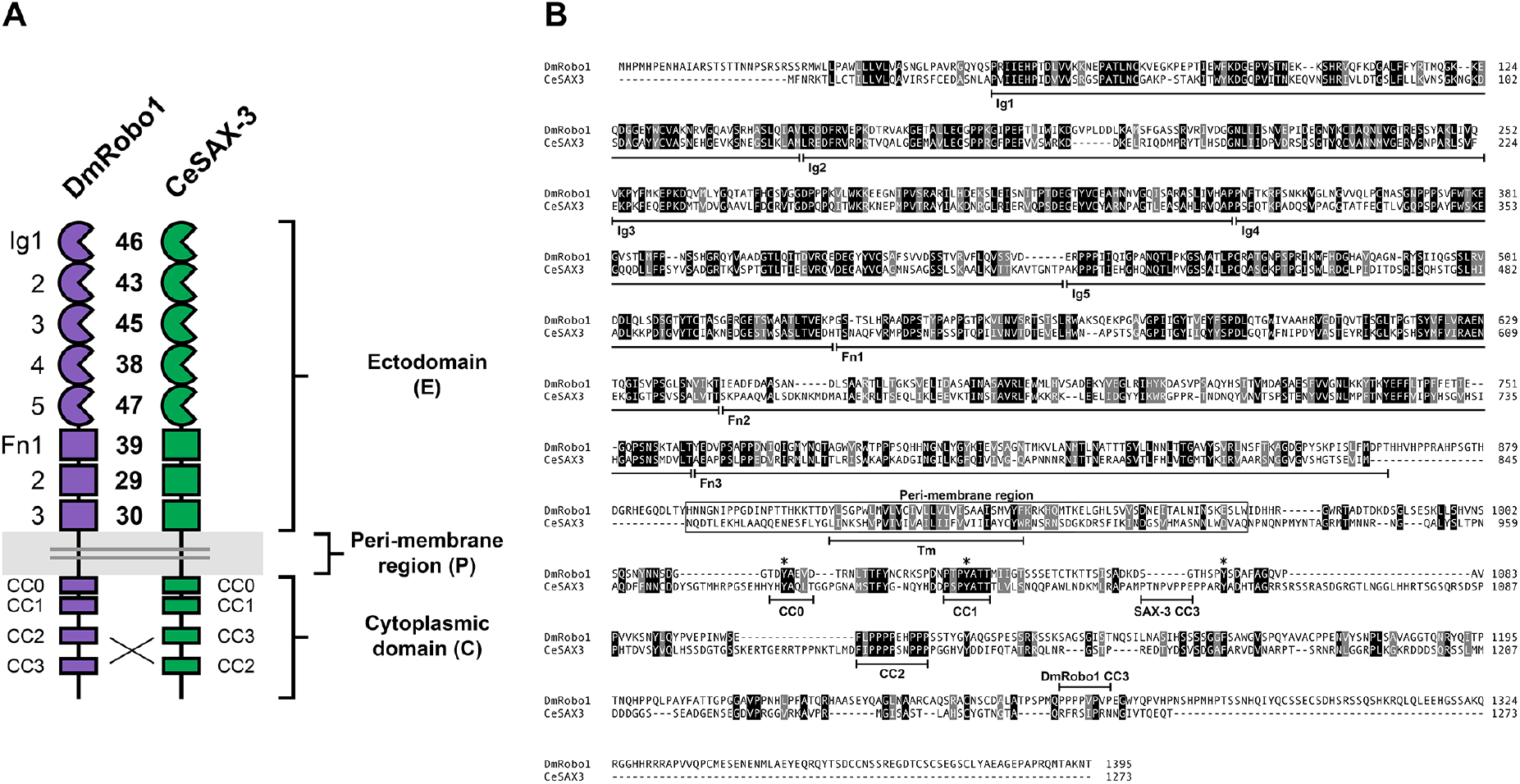
Sequence comparison of *D. melanogaster* Robo1 and *C. elegans* SAX-3. (A) Schematic comparison of the two receptors. Each exhibits the conserved Robo family ectodomain structure including five immunoglobulin-like (Ig) domains and three fibronectin type III (Fn) repeats, and four conserved cytoplasmic (CC) motifs. Numbers indicate percent amino acid identity between the two proteins for each domain. Brackets indicate the extent of ectodomain (E), peri-membrane region (P), and cytoplasmic domain (C). Peri-membrane region is also highlighted in gray. (B) Protein sequence alignment. Structural features are indicated below the sequence. Fn domains have been reannotated relative to Kidd et al. (Kidd *et al*. 1998a) based on revised predictions of beta strand locations. Identical residues are shaded black; similar residues are shaded gray. Ig, immunoglobulin-like domain; Fn, fibronectin type III repeat; Tm, transmembrane helix; CC, conserved cytoplasmic motif. Peri-membrane region (Gilestro 2008) is boxed. Although all four CC motifs (CC0-CC3) are present in both proteins, CC3 is located upstream of CC2 in SAX-3 and thus does not align with Robo1 CC3 (Kidd *et al*. 1998a). Asterisks indicate conserved cytoplasmic tyrosine residues that are phosphorylated by Abl tyrosine kinase in human Robo1 (Bashaw *et al*. 2000).

Although there is much less sequence conservation in the receptors’ cytoplasmic domains, a few areas of strong evolutionary conservation are apparent, including the previously identified CC1, CC2, and CC3 motifs (Fig. 1B). As previously described, the order of motifs is not the same between the two species, with CC3 located upstream of CC2 in SAX-3 but downstream of CC2 in *Drosophila* Robo1 (Kidd *et al*. 1998a). In addition, a short sequence region upstream of CC1 in *C. elegans* SAX-3 (YH<u>YA</u>Q<u>L</u>) includes a tyrosine, alanine, and hydrophobic valine/leucine combination (YAxΦ) that is conserved in the CC0 motif in all three *Drosophila* Robos (Simpson *et al*. 2000b; Rajagopalan *et al*. 2000b; Dickson and Gilestro 2006), leading us to designate this sequence as SAX-3 CC0 (Fig. 1B). Finally, we note that in addition to the conserved tyrosine residues within CC0 and CC1, a third tyrosine position conserved between *Drosophila* Robo1 (HSP<u>Y</u>SDA) and human Robo1 (PVQ<u>Y</u>NIV) and shown to be an Abl phosphorylation target in vitro (Bashaw *et al*. 2000) also appears to be present in the SAX-3 cytodomain (PAR<u>Y</u>ADH), closely adjacent to the CC3 motif (Fig. 1B). We therefore conclude that all four CC motifs (CC0-CC3) and three known phosphorylation sites are present in the cytoplasmic domain of *C. elegans* SAX-3. This conservation suggests that the signaling mechanism(s) downstream of Robo1 and SAX-3 might also be evolutionarily conserved, or at least that SAX-3 might be able to activate downstream components in fly neurons to influence axon guidance and/or signal midline repulsion.

### *C. elegans* SAX-3 can inhibit midline crossing of *Drosophila* axons *in vivo*

To test whether *C. elegans* SAX-3 can regulate axon guidance of *Drosophila* neurons, we first examined the effects of misexpressing SAX-3 broadly in all *Drosophila* neurons using the GAL4/UAS system (Brand and Perrimon 1993). We created a transgenic line of flies carrying a GAL4-responsive *UAS-SAX3* transgene and crossed these flies to a second line carrying a GAL4 transgene expressed in all neurons (*elav-GAL4*). We collected embryos carrying both transgenes (*elav-GAL4/UAS-SAX3*) and examined expression of the transgenic SAX-3 protein using an antibody against an N-terminal HA epitope tag. We examined the effect of SAX-3 misexpression on axon guidance using antibodies which detect all axons in the embryonic CNS (anti-HRP) and a subset of longitudinal axon fascicles (anti-FasII). To compare SAX-3’s activity with that of *Drosophila* Robo1, we performed the same assay using a *UAS-Robo1* transgene (Evans and Bashaw 2012; Brown *et al*. 2015).

In *elav-GAL4/UAS-SAX3* and *elav-GAL4/UAS-Robo1* embryos, transgenic SAX-3 or Robo1 proteins were expressed at similar levels, and both were properly localized to axons in the embryonic ventral nerve cord (Fig. 2D,E). In both misexpression backgrounds, commissural axon tracts were thin or absent in many segments, reflecting ectopic midline repulsion and consistent with our previous analyses of Robo1 misexpression (Fig. 2B,C) (Brown *et al*. 2015). We also observed ectopic midline crossing in some segments in *elav-GAL4/UAS-SAX3* embryos, indicating that SAX-3 misexpression can both promote and inhibit midline crossing when expressed broadly in *Drosophila* embryonic neurons (Fig. 2C). Consistent with this, while transgenic Robo1 protein was only detectable on non-midline-crossing axons, we observed SAX-3 protein at equivalent levels on both midline-crossing and non-crossing axons (Fig. 2E). These results demonstrate that SAX-3 can be properly expressed and localized to axons in *Drosophila* embryonic neurons, and is capable of influencing midline crossing when expressed broadly in all neurons.

**Figure 2.**
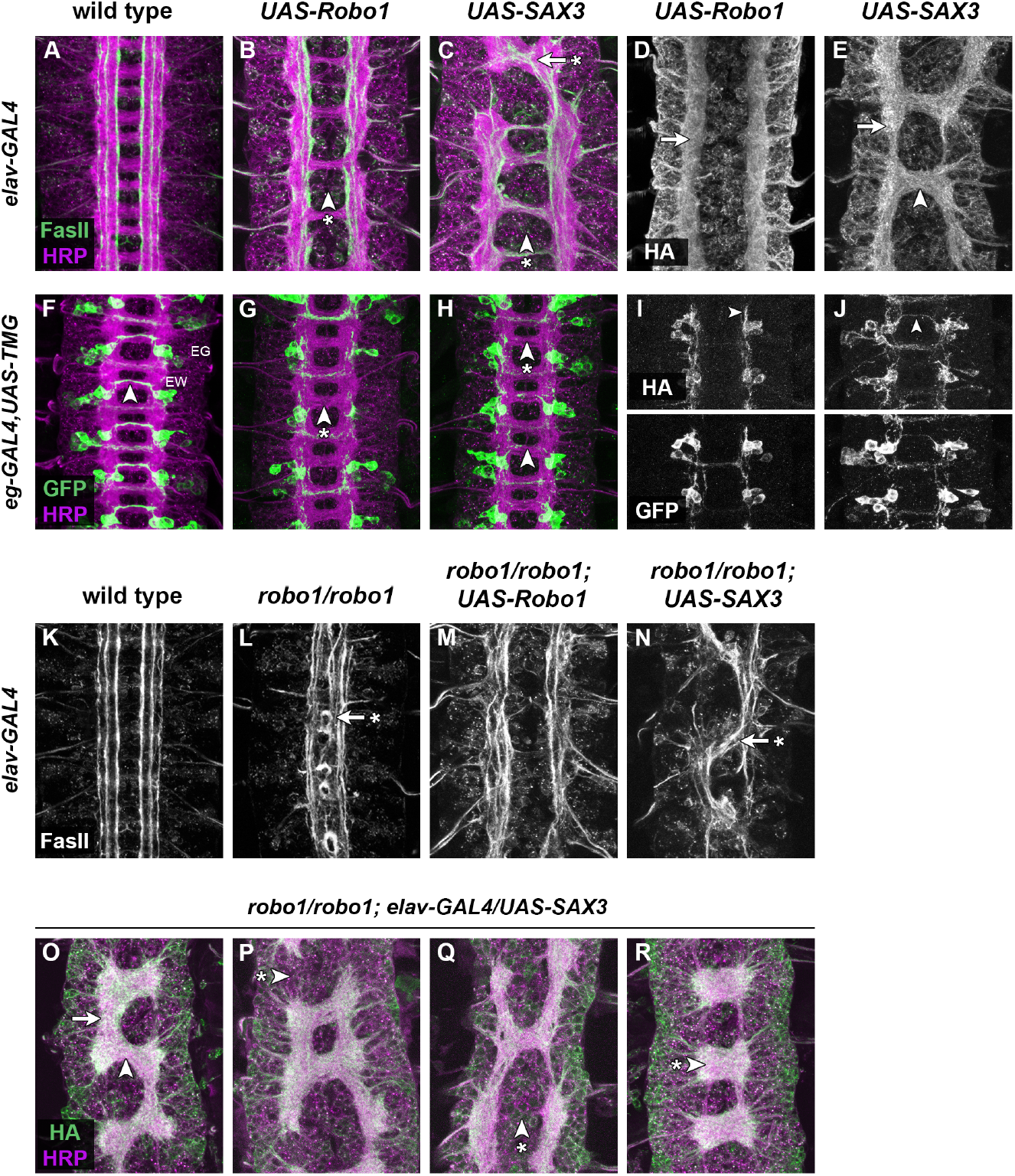
Transgenic SAX-3 can signal midline repulsion in *Drosophila* neurons, but cannot rescue midline repulsion in *robo1* mutants. (A-E) Stage 16 *Drosophila* embryos carrying *elav-GAL4* and the indicated HA-tagged *UAS-Robo* or *UAS-SAX3* transgenes, stained with anti-HRP (magenta; labels all axons) and anti-FasII (green; labels a subset of longitudinal axon pathways) (A-C), or anti-HA (D,E). (A) Embryos carrying *elav-GAL4* alone display a wild type ventral nerve cord with a ladder-like axon scaffold, two commissures per segment, and three distinct FasII-positive longitudinal pathways on either side of the midline. (B) Misexpression of Robo1 with *elav-GAL4* inhibits midline crossing, and commissures are thin or absent (arrowhead with asterisk). (C) In embryos misexpressing SAX-3 with *elav-GAL4*, some segments have reduced or absent commissures (arrowhead with asterisk), while others exhibit thickened or fused commissures with FasII-positive axons ectopically crossing the midline (arrow with asterisk). FasII-positive longitudinal pathways also appear disorganized in (B) and (C). (D,E) Transgenic Robo1 and SAX-3 are both expressed on longitudinal axons (arrows). SAX-3 protein is also detectable on midline-crossing axons (arrowhead in E). (F-J) Stage 15 embryos carrying *eg-GAL4, UAS-TauMycGFP (TMG)*, and the indicated HA-tagged *UAS-Robo* or *UAS-SAX3* transgenes, stained with anti-HRP (magenta) and anti-GFP (green) (F-H), or anti-HA and anti-GFP (I,J). (F) *eg-GAL4* labels the EG and EW neurons, whose axons cross the midline in the anterior and posterior commissures, respectively. In wild type embryos, the EW axons cross the midline in every segment (arrowhead). (G) Misexpression of Robo1 with *eg-GAL4* prevents the EW axons from crossing the midline (arrowhead with asterisk). (H) Misexpression of SAX-3 can also prevent EW axons from crossing the midline (arrowhead with asterisk), but with lower frequency than Robo1. EW axons cross the midline normally in a majority of segments in *eg-GAL4/UAS-SAX3* embryos (arrowhead in H). Anti-HA staining in (I) and (J) shows that both transgenes are expressed on EW axons (arrowheads); GFP staining of the same segments is shown below for comparison. (K-N) Stage 16 embryos stained with anti-FasII. (K) Wild-type embryo carrying *elav-GAL4* alone, as in (A). (L) In *robo1^1^* homozygous null mutant embryos, FasII-positive axons cross the midline ectopically in every segment (arrow with asterisk). Restoring *robo1* expression in all neurons with *elav-GAL4* and *UAS-Robo1* rescues *robo1-* dependent midline repulsion, and FasII-positive axons no longer cross the midline (M). In *robo1* mutant embryos carrying *elav-GAL4* and *UAS-SAX3* (N), FasII-positive axons collapse into one disorganized bundle in most segments, and this bundle frequently crosses the midline (arrow with asterisk). (O-R) Stage 16 embryos stained with anti-HRP (magenta) and anti-HA (green). In *robo1* mutants carrying *elav-GAL4* and *UAS-SAX3*, the HA-tagged transgenic SAX-3 protein is detectable on both longitudinal connectives (O, arrow) and commissures (O, arrowhead). These embryos display a combination of breaks in the longitudinal connectives (arrowhead with asterisk in P), thin or absent commissures (arrowhead with asterisk in Q), and fused commissures (arrowhead with asterisk in R).

To more closely examine the ability of SAX-3 to repel axons from the midline in the *Drosophila* embryonic CNS, we used *eg-GAL4*, a more restricted GAL4 line that is expressed in two distinct subsets of commissural neurons (the EG and EW neurons). In this experiment we also included a *UAS-TauMycGFP* (*UAS-TMG*) transgene to label the EG and EW cell bodies and axons with GFP. While EW axons cross the midline in 100% of segments in *eg-GAL4,UAS-TMG* control embryos, Robo1 or SAX-3 misexpression was capable of preventing EW axon crossing. We found that misexpression of SAX-3 with *eg-GAL4* prevented midline crossing of EW axons in 42.5% of segments (n=118 segments in 15 embryos), compared to equivalent expression of Robo1 which prevented EW axons from crossing in 96.9% of segments (n=96 segments in 12 embryos) (Fig. 2F-H). We therefore conclude that *C. elegans* SAX-3 can activate midline repulsive signaling in *Drosophila* embryonic neurons and can prevent midline crossing when expressed broadly or in a restricted subset of commissural neurons.

### Pan-neural expression of SAX-3 is unable to rescue midline crossing in *robo1* mutants

The gain-of-function experiments described above demonstrate that SAX-3 can induce ectopic midline repulsion when expressed in *Drosophila* neurons. Importantly, these experiments were carried out in embryos expressing normal levels of endogenous Robo1. To determine whether SAX-3 can promote midline repulsion in embryos lacking *robo1*, we performed a rescue assay using our *UAS-Robo1* and *UAS-SAX3* transgenes in *robo1* null mutants (Brown *et al*. 2015). In *robo1^1^* homozygous mutant embryos, FasII-positive axons cross the midline in 100% of segments; these axons do not cross the midline in wild type embryos (Fig. 2K,L). Forcing high levels of Robo1 expression in all neurons in *robo1* mutants restores midline repulsion and rescues the ectopic midline crossing phenotype, although expressing Robo1 at such high levels in *robo1* mutants also induces additional defects including ectopic midline repulsion and disorganization of longitudinal axon pathways (Fig. 2M) (Brown *et al*. 2015). When we forced high-level SAX-3 expression in all neurons in *robo1* mutant embryos (*robo1^1^/robo1^1^; elav-GAL4/UAS-SAX3*), we observed a number of severe defects that were distinct from the stereotypical midline crossing defects seen in *robo1* mutants (Fig. 2N, O-R). In many segments, all of the longitudinal axons collapsed together and crossed the midline together, resulting in longitudinal breaks or gaps in the axon scaffold. Some segments lacked commissures completely, while in others a single thick commissural bundle was present. In most segments axons did not linger at the midline or re-cross within the same segment, suggesting that some level of midline repulsion is intact in these embryos. However, the additional defects caused by high levels of SAX-3 misexpression in *robo1* mutant embryos prevented us from accurately measuring the extent to which SAX-3 could substitute for Robo1 to signal midline repulsion (if any).

### Expression of SAX-3 in *Drosophila* embryonic neurons via a *robo1* rescue transgene

To more accurately compare the midline repulsive activity of SAX-3 and Robo1 in *Drosophila* neurons, we next used a *robo1* rescue transgene that includes regulatory sequences from the *Drosophila robo1* gene to express *sax-3* in a pattern and expression level that closely reproduces the endogenous expression of *robo1* (Fig. 3A). We have previously used this transgenic approach to perform structure-function analyses of Robo1 ectodomain elements (Brown *et al*. 2015; Reichert *et al*. 2016; Brown *et al*. 2018; Brown and Evans 2020). All of the constructs described herein include the endogenous signal peptide from Robo1 and a 4xHA epitope tag inserted directly upstream of the Ig1 domain, and were inserted at the same genomic location (28E7) to ensure equivalent expression levels between transgenes (Fig. 3A).

**Figure 3.**
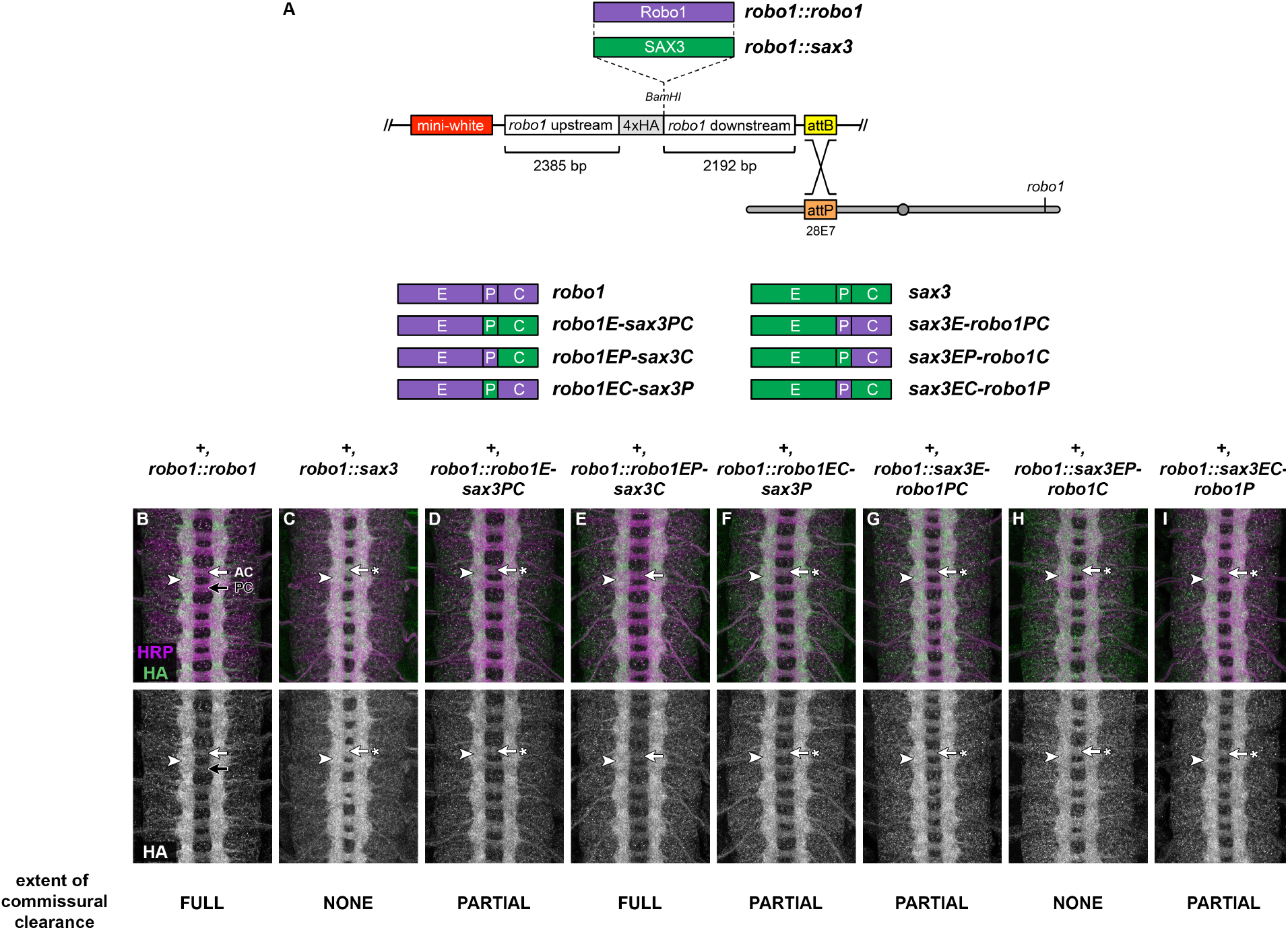
Expression and localization of SAX-3 and Robo1/SAX-3 chimeric receptors via a *robo1* genomic rescue construct. (A) Schematic of the *robo1* rescue construct (Brown *et al*. 2015) and Robo1/SAX-3 chimeric receptors carrying different combinations of ectodomain (E), peri-membrane (P), and cytoplasmic (C) sequences, as defined in Figure 1. See Methods for precise amino acid compositions. HA-tagged receptors are expressed under the control of regulatory regions from the *robo1* gene. All transgenes are inserted into the same genomic landing site at cytological position 28E7. (B-I) Stage 16 embryos stained with anti-HRP (magenta) and anti-HA (green) antibodies. Bottom images show HA channel alone from the same embryos. HA-tagged full-length Robo1 (B) expressed from the *robo1* rescue transgene is localized to longitudinal axon pathways (arrowhead) and excluded from commissural segments in both the anterior commissure (AC, white arrow) and posterior commissure (PC, black arrow). (B) Transgenic SAX-3 protein expressed from an equivalent transgene is properly localized to axons (arrowhead), but is not excluded from commissures (arrow with asterisk). Each of the Robo1/SAX-3 chimeric receptors is also properly localized to axons (arrowheads in D-I), but display varying degrees of commissural exclusion, from nearly complete exclusion similar to full-length Robo1 (*robo1EP-sax3c*, E, arrow), to partial exclusion (*robo1E-sax3PC*, D; *robo1EC-sax3P*, F; *sax3EC-robo1P*, I, arrows with asterisk), to no exclusion similar to full-length SAX-3 (*sax3EP-robo1C*, H, arrow with asterisk).

As we have previously described, expression of full-length transgenic Robo1 from our rescue construct accurately reproduced the endogenous expression pattern of Robo1, with the HA-tagged transgenic Robo1 protein detectable on longitudinal axons in the embryonic ventral nerve cord but largely excluded from commissural axon segments (Fig. 3B). We found that SAX-3 protein expressed from an equivalent transgene was also properly translated, expressed at similar levels to Robo1, and localized to axons in the embryonic CNS, but unlike Robo1 was not excluded from commissures (Fig. 3C). Homozygous embryos carrying two copies of our *robo1::sax3* transgene in addition to two wild-type copies of the endogenous *robo1* gene (*+, robo1::sax3*) displayed slightly thickened commissures and a reduced distance between the longitudinal connectives and the midline, suggesting a mild dominant-negative effect caused by SAX-3 expression in otherwise wild-type embryos (Fig. 3C). Staining with an anti-FasII antibody confirmed a low level of ectopic midline crossing in these embryos (not shown).

### SAX-3 can partially rescue midline repulsion in the absence of *robo1*

To determine whether SAX-3 can substitute for Robo1 to promote midline repulsion during embryonic development, we introduced our *robo1::sax3* transgene into a *robo1* null mutant background and examined midline repulsion using anti-FasII, which labels a subset of longitudinal axon pathways in the *Drosophila* embryonic CNS.

In wild type stage 16-17 *Drosophila* embryos, FasII-positive axons do not cross the midline (Fig. 4A), while in *robo1* null mutants they cross the midline ectopically in 100% of segments in the ventral nerve cord (Fig. 4B). Restoring expression of full-length Robo1 via the *robo1::robo1* rescue transgene completely rescues midline repulsion in *robo1* mutant embryos (Fig. 4C) (Brown *et al*. 2015). Expressing *sax-3* in *robo1’s* normal pattern partially restored midline repulsion in embryos lacking endogenous *robo1* (*robo1^1^, robo1::sax3*) (Fig. 4D). In these embryos, FasII-positive axons crossed the midline in 64.8% of segments, which represents a significant rescue compared to *robo1* null mutants (p<0.001 by Student’s t-test). These results demonstrate that SAX-3 can signal midline repulsion in *Drosophila* embryonic neurons and can substitute for *Drosophila robo1* in its endogenous context of preventing longitudinal axons from crossing the midline, although it cannot perform this role as effectively as *robo1*.

**Figure 4.**
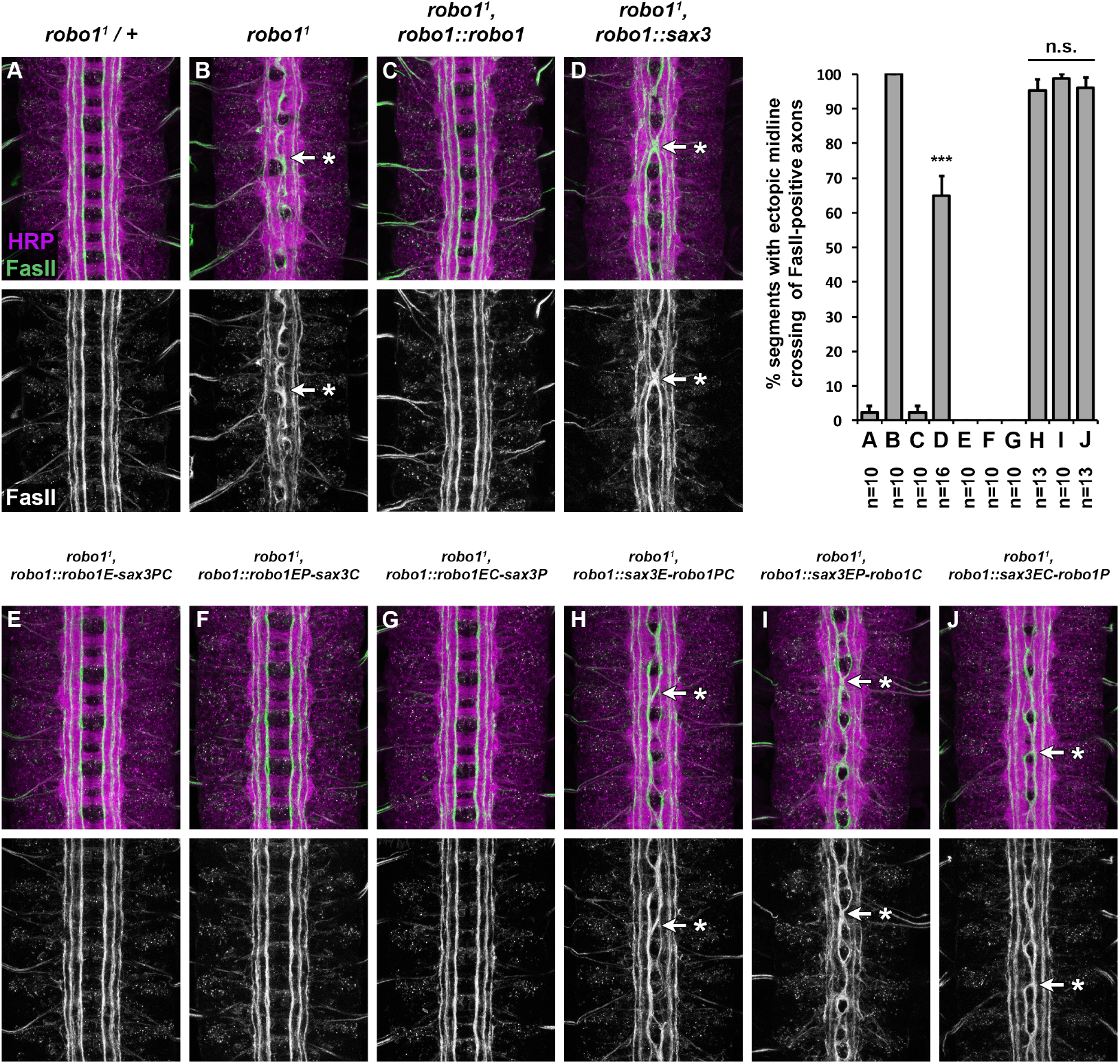
Full-length SAX-3 and Robo1/SAX-3 chimera rescue. (A-J) Stage 16 *Drosophila* embryos stained with anti-HRP (magenta) and anti-FasII (green) antibodies. Lower images show anti-FasII channel alone from the same embryos. FasII-positive axons cross the midline ectopically in *robo1* mutant embryos (B, arrow with asterisk), and this defect is rescued completely by restoring Robo1 expression via a genomic rescue transgene (C). Expression of SAX-3 in *robo1’s* normal expression pattern partially rescues midline repulsion, and FasII-positive axons cross the midline in 64.8% of segments in these embryos (D). (E-J) *robo1* mutant embryos expressing Robo1/SAX-3 chimeric receptors via the *robo1* rescue transgene. Midline crossing defects are completely rescued in embryos expressing chimeric receptors that contain the ectodomain of Robo1 (*robo1E-sax3PC, E; robo1EP-sax3C, F; robo1EC-sax3P, G*) but not rescued by chimeric receptors that contain the SAX-3 ectodomain (*sax3E-robo1PC, H; sax3EP-robo1C, I; sax3EC-robo1P, J;* arrows with asterisks). Bar graph shows quantification of ectopic midline crossing defects in the genotypes shown in (A-J). Error bars indicate standard error. Number of embryos scored (n) is shown for each genotype. Full-length SAX-3 and Robo1/SAX-3 chimeric receptor rescue phenotypes (D,H-J) were compared to *robo1* mutant embryos (B) by Student’s t-test, with a Bonferroni correction for multiple comparisons (***p<0.0001).

### Robo1/SAX-3 chimeric receptor variants

We have noted two differences between Robo1 and SAX-3 when they are expressed in *Drosophila* embryonic neurons: first, Robo1 protein is largely restricted to longitudinal axons and excluded from commissures, while SAX-3 is expressed uniformly on both longitudinal and commissural axon segments; second, transgenic Robo1 is able to fully rescue midline crossing defects in a *robo1* mutant background, while SAX-3 can only partially rescue these defects. We hypothesized that differences in axonal distribution might be due to sequence differences in the fibronectin repeats (Fn) or peri-membrane region, which have been implicated in commissural clearance and/or regulation of *Drosophila* Robo1 by Commissureless, which prevents surface localization of Robo1 protein in pre-crossing commissural neurons (Gilestro 2008; Brown *et al*. 2018). Further, we suspected that SAX-3’s reduced ability to rescue midline repulsion might be due to differences in binding or responding to *Drosophila* Slit, which should be ectodomain-dependent, or downstream signaling output, which would be cytodomain-dependent.

To address these possibilities, we constructed a series of chimeric receptors combining the ectodomain (E), peri-membrane (P), and cytoplasmic (C) regions of Robo1 and SAX-3 (Fig. 1A, Fig. 3A). We defined the ectodomains (E) of Robo1 and SAX-3 as all sequences upstream of the 83 aa peri-membrane region of Robo1 defined by Gilestro (amino acids H891–W973) (Gilestro 2008) or the equivalent region of SAX-3 (amino acids N847–Q949), while the cytoplasmic domains (C) were defined as all sequences downstream of this region. By these definitions, all ectodomain structural elements (Ig1-5, Fn1-3) are included in the ectodomain (E) region, while all of the conserved cytoplasmic (CC) motifs are included in the cytoplasmic (C) domain. The peri-membrane (P) region includes the transmembrane helix in addition to 21 aa upstream and 33 aa downstream in both proteins (Fig. 1A).

We then made transgenic lines for each chimeric receptor using our *robo1* rescue transgene construct, and we examined the expression and localization of these HA-tagged chimeric receptors in *Drosophila* embryonic neurons, as well as their ability to rescue midline crossing defects in a *robo1* null mutant background.

### Robo1 ectodomain and perimembrane regions both contribute to commissural clearance

When expressed from our *robo1* rescue construct, transgenic Robo1 protein is present on longitudinal axons but largely absent from commissures, while transgenic SAX-3 is uniformly expressed on both longitudinal and commissural axon segments. We found that our Robo1/SAX-3 chimeric receptors were all properly translated and localized to axons, and displayed varying levels of commissural clearance which correlated with the presence of the Robo1 ectodomain and peri-membrane regions. The variant that included both ectodomain (E) and peri-membrane (P) from Robo1 (robo1EP-sax3C) displayed nearly complete commissural clearance similar to full-length Robo1 (Fig. 3E), while the variant that included both ectodomain (E) and peri-membrane (P) from SAX-3 (sax3EP-robo1C) was not cleared at all, similar to fulllength SAX-3 (Fig. 3H). The variants that included either the ectodomain (E) or peri-membrane (P) from Robo1, but not both (robo1E-sax3PC, robo1EC-sax3P, sax3E-robo1PC, sax3EC-robo1P), displayed intermediate levels of commissural clearance (Fig. 3 D,F,G,I). These results suggest that both the ectodomain and peri-membrane regions of Robo1 contribute to its exclusion from commissural axon segments, and that their contributions to commissural exclusion may be additive. This is consistent with our previously described Robo1ΔFn3 variant, which includes an intact peri-membrane region and also displays partial clearance from commissures (Brown *et al*. 2018), and suggests that whatever sequence(s) within Robo1 Fn3 contribute to commissural clearance are not conserved in SAX-3.

### Chimeric Robo1/SAX-3 receptors containing the Robo1 ectodomain can fully rescue midline repulsion in *robo1* mutants

We next asked whether our Robo1/SAX-3 chimeric receptors could rescue midline crossing defects caused by loss of *robo1*. To this end, as with our full-length *robo1::sax3* transgene described above, we introduced each transgene into a *robo1* null mutant background and quantified ectopic crossing of FasII-positive longitudinal axons in stage 16-17 embryos (Fig. 4E-J). We found that each of the receptor variants that included Robo1’s ectodomain (robo1E-sax3PC, robo1EP-sax3C, robo1EC-sax3P) could fully rescue midline crossing defects in *robo1* mutants, equivalent to full-length Robo1 (Fig. 4E-G), while variants that included SAX-3’s ectodomain (sax3E-robo1PC, sax3EP-robo1C, sax3EC-robo1P) could not rescue midline repulsion (Fig. 4H-J). These results suggest that SAX-3’s reduced ability to rescue *robo1*-dependent midline repulsion is due to functional difference(s) between the Robo1 and SAX-3 ectodomains, perhaps their relative affinities for *Drosophila* Slit. These results also demonstrate that the SAX-3 cytoplasmic domain is able to activate downstream repulsive signaling in *Drosophila* neurons as effectively as the Robo1 cytoplasmic domain, as long as it is paired with Robo1’s ectodomain.

### Robo1 ectodomain and perimembrane regions are both required for down-regulation by Comm

Commissureless (Comm) is a negative regulator of Slit-Robo signaling in *Drosophila* that prevents newly-synthesized Robo1 protein from reaching the growth cone surface in precrossing commissural axons (Tear *et al*. 1996; Kidd *et al*. 1998b; Keleman *et al*. 2002; 2005; Gilestro 2008). *comm* is normally expressed transiently in commissural neurons as their axons are crossing the midline, and its transcription is rapidly extinguished after midline crossing (Keleman *et al*. 2002). Accordingly, forced expression of Comm in all neurons leads to a strong reduction in Robo1 protein levels and an ectopic midline crossing phenotype that resembles *robo1* or *slit* loss of function mutants (Kidd *et al*. 1998b; Gilestro 2008; Brown *et al*. 2015; Reichert *et al*. 2016; Brown *et al*. 2018). Notably, Robo1 variants that are resistant to endosomal sorting by Comm can still be antagonized by Comm via an uncharacterized but sorting-independent mechanism (Gilestro 2008). We and others have shown that the perimembrane region and Fn3 domain of Robo1 are each required for downregulation by Comm, but Robo1 variants lacking these regions are still subject to sorting-independent Comm antagonism (Gilestro 2008; Brown *et al*. 2018). Comm appears to be conserved only within insects, and Comm orthologs have not been identified in nematodes or other animal groups (though Ndfip and PRRG4 proteins have been proposed as functional analogs of Comm in mammals (Justice *et al*. 2017; Gorla *et al*. 2019)). We therefore asked whether SAX-3 is sensitive to endosomal sorting or sorting-independent antagonism by *Drosophila* Comm when expressed in *Drosophila* neurons. We used *elav-GAL4* and *UAS-Comm* transgenes to drive high levels of Comm expression in all neurons in embryos carrying one copy of our Robo1, SAX-3, or Robo1/SAX-3 chimeric receptor transgenes, then examined the effect on the expression levels of each transgene along with axon scaffold architecture using anti-HA and anti-HRP antibodies (Fig. 5).

**Figure 5.**
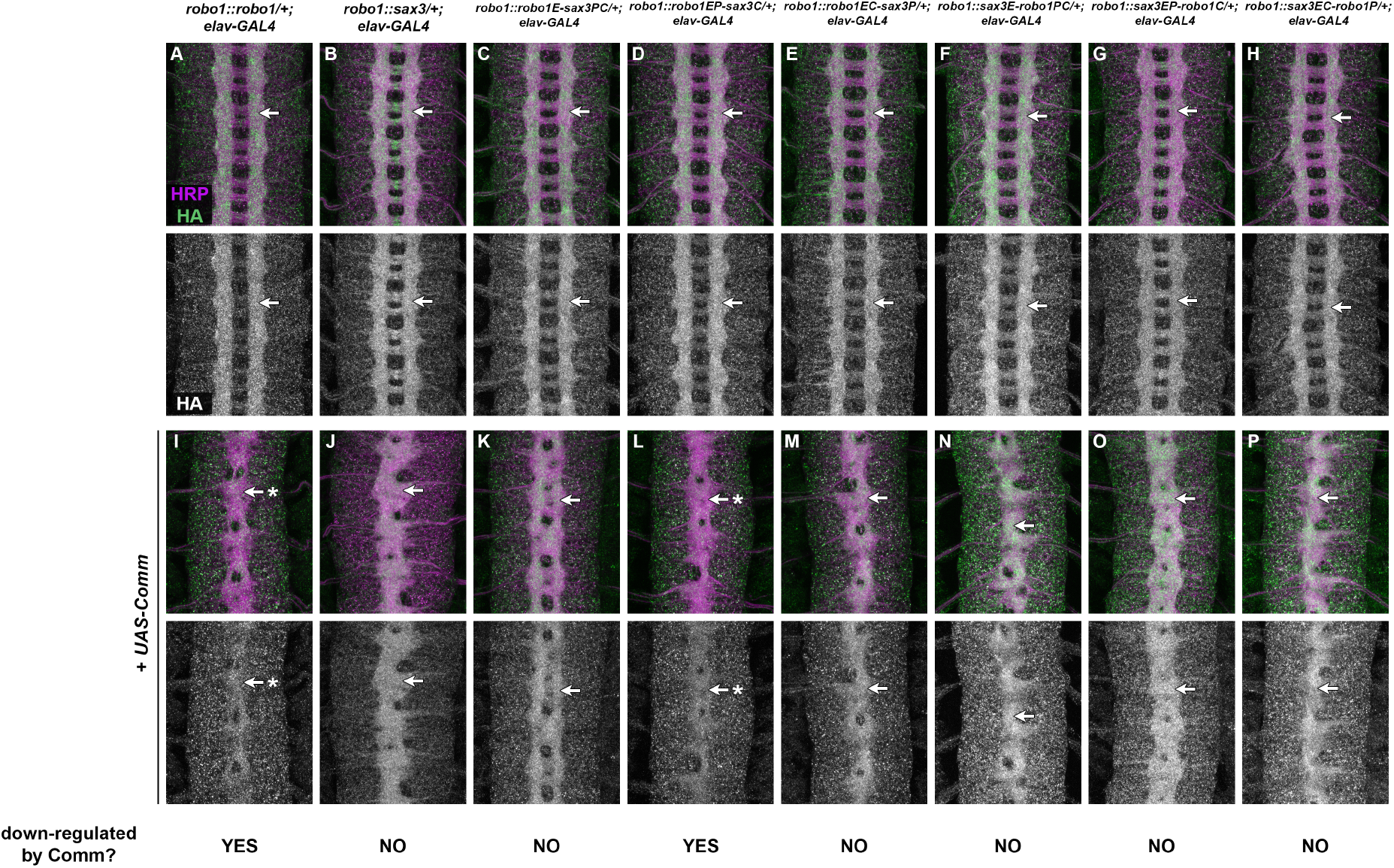
Comm-dependent downregulation of Robo1/SAX-3 chimeras. (A-P) Stage 16 *Drosophila* embryos carrying one copy of the indicated *robo1* rescue transgenes along with *elav-GAL4* alone (A-H) or *elav-GAL4* and *UAS-Comm* (I-P). All embryos are stained with anti-HRP (magenta) to label the entire axon scaffold and anti-HA (green) to detect expression level of transgenic Robo proteins. All embryos also carry two wild-type copies of the endogenous *robo1* gene. Embryos carrying one copy of the indicated *robo1* transgenes along with *elav-GAL4* display normal expression of the HA tagged transgenic receptor variants (A-H, arrows). Expression levels of full-length Robo1 (I) and Robo1EP-SAX3C (L) are strongly decreased when *comm* is expressed at high levels in all neurons (I,L, arrows with asterisk), but expression levels of the other chimeric receptors or full-length SAX-3 are unaffected by ectopic *comm* expression (J,K,M-P, arrows). In all backgrounds, *comm* misexpression causes ectopic midline crossing resulting in a *robo1*-like or *slit*-like midline collapse phenotype, as revealed by anti-HRP (I-P). Pairs of sibling embryos shown here (A and I; B and J; C and K; D and L; E and M; F and N; G and O; H and P) were stained in the same tube and imaged using identical confocal settings to allow accurate comparisons of HA levels between embryos with and without ectopic *comm* expression.

As we have previously described, Comm misexpression leads to a strong reduction in transgenic Robo1 protein as well as midline collapse of the axon scaffold, compared to sibling embryos carrying *elav-GAL4* only, reflecting endosomal sorting and degradation of Robo1 protein in the presence of Comm (Fig. 5A,I) (Brown *et al*. 2015; Reichert *et al*. 2016; Brown *et al*. 2018). In contrast, we found that Comm misexpression produces a strong midline collapse phenotype without altering SAX-3 levels in embryos carrying our *robo1::sax3* transgene (Fig. 5B,J). This suggests that SAX-3 is insensitive to endosomal sorting by Comm, but still subject to sorting-independent antagonism. Among our Robo1/SAX-3 chimeric receptor transgenes, all were insensitive to Comm sorting except the chimeric receptor that included the ectodomain and peri-membrane region of Robo1 (robo1EP-sax3C) (Fig. 5D,L), supporting the idea that both Fn3 and peri-membrane regions are required for endosomal sorting of Robo1 by Comm *in vivo*. Like full-length Robo1 and SAX-3, all of the chimeric receptor transgenes were sensitive to sorting-independent antagonism by Comm, as evidenced by the midline collapse phenotype caused by Comm misexpression in embryos carrying any of our rescue transgenes (Fig. 5I-P).

### SAX-3 dependent midline repulsion in Drosophila neurons is hyperactive in *comm* mutants

The above results demonstrate that *Drosophila* Comm is unable to downregulate *C. elegans* SAX-3 expression in *Drosophila* neurons, but SAX-3 is still sensitive to sorting-independent regulation by Comm when Comm is misexpressed in all neurons. In addition, embryos carrying the *robo1::sax3* transgene are not commissureless (see Fig. 3C and 4D), suggesting that Comm may be regulating SAX-3-mediated midline repulsion in these embryos. Alternatively, SAX-3 may be completely free of Comm inhibition in these embryos, and the lack of a commissureless phenotype in *robo1::sax3* embryos may be due solely to the fact that SAX-3 cannot signal midline repulsion as efficiently as Robo1 in *Drosophila* neurons.

To distinguish between these possibilities, we examined the effect of removing *comm* in embryos expressing either the *robo1::robo1* or *robo1::sax3* transgenes in place of endogenous *robo1*. Amorphic *comm* mutant embryos display a strongly commissureless phenotype, with few or no axons crossing the midline (Fig. 6B). This phenotype is *robo1-*dependent, as *robo1,comm* double mutant embryos phenocopy *robo1* mutants (Seeger *et al*. 1993). Thus, the commissureless phenotype seen in *comm* embryos is due to hyperactivity of endogenous *robo1. robo1, comm* double mutants carrying our *robo1::robo1* rescue transgene also display a commissureless phenotype, indicating that transgenic Robo1 expressed from the rescue transgene is hyperactive in the absence of *comm* (Fig. 6C). Similarly, *robo1,comm* double mutants carrying our *robo1::sax3* rescue transgene also display a strongly commissureless phenotype, although one that is slightly less severe than *comm* mutants alone or *robo1,robo1::robo1; comm* compound mutants (Fig. 6D). We infer from this phenotype that SAX-3 protein expressed from the *robo1::sax3* transgene is normally subject to negative regulation by *comm*, and SAX-3-dependent midline repulsive signaling becomes hyperactive in the absence of *comm*. The slight difference is severity of the commisureless phenotypes in *robo1,robo1::robo1; comm* and *robo1,robo1::sax3; comm* embryos likely reflects the quantitative difference in signaling activity of Robo1 and SAX-3 in *Drosophila* neurons.

**Figure 6.**
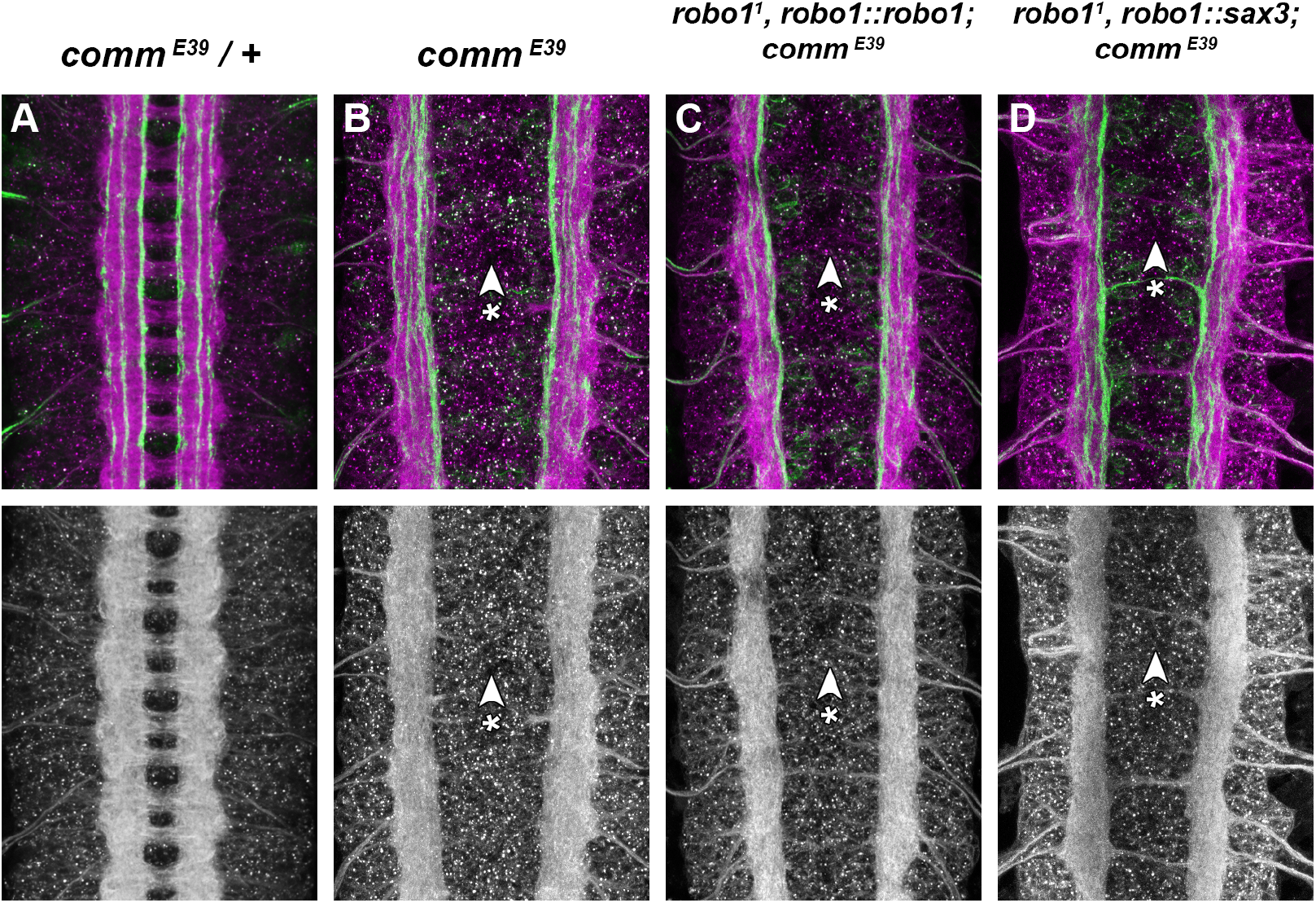
SAX-3-mediated midline repulsion is hyperactive in the absence of *comm*. (A-D) Stage 16 *Drosophila* embryos stained with anti-HRP (magenta) and anti-FasII (green) antibodies. Lower images show anti-HRP channel alone from the same embryos. The axon scaffold forms normally in embryos heterozygous for an amorphic allele of *comm* (A). In homozygous *comm* mutant embryos, commissural axons fail to cross the midline and the commissures do not form because endogenous Robo1 is hyperactive (B, arrowhead with asterisk). This phenotype is reproduced in *robo1,comm* double mutant embryos carrying the *robo1::robo1* transgene, indicating that the Robo1 protein expressed from the transgene is hyperactive in the absence of *comm* (C, arrowhead with asterisk). *robo1,comm* double mutants carrying the *robo1::sax-3* transgene also display strongly reduced commissures, indicating that SAX-3 is also hyperactive in embryos lacking *comm* (D, arrowhead with asterisk).

## Discussion

In this paper, we have used a transgenic approach in the *Drosophila* embryonic CNS to examine the evolutionary conservation of midline repulsive signaling activity between two members of the Roundabout (Robo) family of axon guidance receptors: *Drosophila melanogaster* Robo1 and *Caenorhabditis elegans* SAX-3. Robo1 and SAX-3 were two of the first Robo family members to be described, and their similar protein structure and developmental roles suggested strong evolutionary conservation of midline repulsive signaling mechanisms across animal groups (Kidd *et al*. 1998a; Zallen *et al*. 1998). Here, we have directly examined this functional conservation by expressing SAX-3 in *Drosophila* embryonic neurons and testing its ability to signal midline repulsion as well as to substitute for *Drosophila robo1* to regulate midline crossing during embryonic development. We show that *C. elegans* SAX-3 can prevent *Drosophila* axons from crossing the midline, presumably by signaling midline repulsion in response to *Drosophila* Slit, and can partially substitute for *Drosophila robo1* to properly regulate midline crossing of longitudinal axons in the *Drosophila* embryonic CNS. Using a series of chimeric receptors, we show that the SAX-3 cytoplasmic domain can act equivalently to the Robo1 cytoplasmic domain to signal midline repulsion in *Drosophila* neurons when combined with the Robo1 ectodomain, but reciprocal chimeras combining the SAX-3 ectodomain with the Robo1 cytodomain cannot effectively signal midline repulsion. We further show that SAX-3 is insensitive to endosomal sorting by *Drosophila* Comm, but is subject to sorting-independent antagonism by Comm.

### The SAX-3 and Robo1 cytodomains can act equivalently to signal midline repulsion in *Drosophila* axons

Our Robo1/SAX-3 chimeric receptors reveal that the SAX-3 cytodomain can signal midline repulsion in *Drosophila* neurons when paired with the ectodomain of Robo1, at a level that is indistinguishable from the native Robo1 cytodomain. While this observation does not directly demonstrate that the signaling mechanisms downstream of Robo1 and SAX-3 are identical in fly and worm neurons, it does suggest that the SAX-3 cytodomain is capable of interacting with and activating the downstream signaling components necessary for midline repulsive signaling in fly neurons. Our sequence comparisons indicate that all four of the previously identified CC motifs (CC0-CC3) are present in the SAX-3 cytodomain, along with a third Abl phosphorylation site (in addition to the two in CC0 and CC1) that is outside of these four motifs. This supports the idea that all critical signaling elements are present in the SAX-3 cytodomain. Further, since the order of the CC2 and CC3 motifs are switched relative to the Robo1 cytodomain, and the spacing between these sequences is not identical in the two receptors, this indicates that the order and relative positions of these sequence elements is not critical for their signaling output. Long and colleagues have used a similar chimeric receptor approach to show that the cytodomains of the unrelated repulsive axon guidance receptors Derailed (Drl) and Unc5 can also substitute for the Robo1 cytodomain to rescue *robo1*-dependent midline repulsion when combined with the Robo1 ectodomain (Long *et al*. 2016), suggesting that all three of these receptors may function through a common downstream signaling pathway. Our results indicate that the SAX-3 cytodomain can also activate this pathway.

### Ectodomain-dependent differences in midline repulsive output

Both the structural ectodomain arrangment of 5 Ig + 3 Fn domains and the Slit-binding sequences within the N-terminal Ig1 domain are highly conserved across Robo receptors in many bilaterian species, while the cytoplasmic domain sequences are much more divergent. Ectodomain-dependent differences in activities between *Drosophila* Robo paralogs have been described in contexts other than midline repulsion (Evans and Bashaw 2010; Evans *et al*. 2015), while differences between *Drosophila* Robo1’s and Robo3’s abilities to rescue midline repulsion in *robo1* mutants has been attributed entirely to differences in their cytoplasmic domains (namely, the CC1-CC2 region of Robo1) (Spitzweck *et al*. 2010). However, the chimeric receptors described here indicate that the cytoplasmic domains of *Drosophila* Robo1 and *C. elegans* SAX-3 are functionally interchangeable in the context of midline repulsion in *Drosophila* neurons, while their ectodomains are not. One possibility is that there are quantitative differences in affinity for *Drosophila* Slit caused by sequence divergence within Ig1 (which is 46% identical between Robo1 and SAX-3; Fig. 1). Although it is clear that SAX-3 can detect and respond to *Drosophila* Slit, it does so less efficiently than *Drosophila* Robo1, as seen in both the gain of function and *robo1* mutant rescue experiments presented here. Another possibility is that, independent of Slit affinity, there may be ectodomain conformational arrangments or changes in response to Slit binding that are necessary for optimal signaling that SAX-3 does not share with Robo1. In either case, it is unclear why chimeras containing the SAX-3 ectodomain (which cannot rescue midline repulsion at all in *robo1* mutants) would be less active than fulllength SAX-3 (which can partially rescue midline repulsion). Perhaps there are quantitative effects of ectodomain/cytodomain compatibility that further reduce the repulsive signaling efficiency when the SAX-3 ectodomain is paired with the Robo1 cytodomain.

### Antagonism of SAX-3-dependent midline repulsion by Comm

Comm is able to antagonize *Drosophila* Robo1 through two apparently distinct mechanisms: one that involves endosomal sorting of Robo1 and depends on both the peri-membrane and Fn3 regions of Robo1 (Keleman *et al*. 2002; 2005; Gilestro 2008; Brown *et al*. 2018), and a second, as yet uncharacterized sorting-independent mechanism (Gilestro 2008). It is unclear whether sorting-independent regulation of Slit-Robo1 repulsion by Comm is achieved through direct regulation of Robo, or through some other component(s) of the Slit-Robo1 pathway (Gilestro 2008). The epistatic relationship between *comm* and *robo1* indicates that this regulation must occur at the level of *robo1* or upstream (Seeger *et al*. 1993). Whatever the mechanism or level of action of this sorting-independent regulation, SAX-3 must also be sensitive to it, as removal of *comm* in *robo1^1^, [robo1::sax3]* embryos produces a commissureless phenotype indicative of hyperactive Slit-Robo repulsion (Fig. 6D). There do not appear to be any *comm* orthologs present in *C. elegans*. Perhaps *comm* regulation does not depend on specific sequences in Robo1/SAX-3 or, alternatively, this regulation may rely on sequences or structural arrangements in Robo1 that are conserved in SAX-3 for other reasons.

## Acknowledgments

We thank Cori Bargmann for providing the *sax-3* cDNA clone, and Nicholas Jones for contributing to the cloning of the *[robo1::sax3]* transgene. Stocks obtained from the Bloomington Drosophila Stock Center [National Institutes of Health (NIH) grant P40 OD-018537] were used in this study. Monoclonal antibodies were obtained from the Developmental Studies Hybridoma Bank, created by the Eunice Kennedy Shriver National Institute of Child Health and Human Development of the NIH and maintained at The Department of Biology, University of Iowa, Iowa City, IA 52242. This work was supported by NIH grant R15 NS-098406 (T.A.E.).

